# *Treponema pallidum* periplasmic and membrane proteins are recognized by circulating and skin CD4+ T cells

**DOI:** 10.1101/2024.02.27.581790

**Authors:** Tara B. Reid, Charmie Godornes, Victoria L. Campbell, Kerry J. Laing, Lauren C. Tantalo, Alloysius Gomez, Thepthara N. Pholsena, Nicole A. P. Lieberman, Taylor M. Krause, Victoria I. Cegielski, Lauren A. Culver, Nhi Nguyen, Denise Q. Tong, Kelly L. Hawley, Alexander L. Greninger, Lorenzo Giacani, Caroline E. Cameron, Julia C. Dombrowski, Anna Wald, David M. Koelle

**Affiliations:** Department of Medicine, University of Washington School of Medicine, Seattle, WA, USA; Department of Biochemistry and Microbiology, University of Victoria, BC, Canada; Department of Laboratory Medicine and Pathology, University of Washington, Seattle, WA, USA; University of Missouri-Kansas City School of Medicine, Kansas City, MO, USA; Department of Medicine and Pediatrics, UConn Health, Farmington, CT, USA; Division of Infectious Diseases, Connecticut Children’s, Hartford CT, US; Vaccine and Infectious Diseases Division, Fred Hutchinson Cancer Center, Seattle, WA, USA; Department of Global Health, University of Washington, Seattle, WA, USA; Department of Epidemiology, University of Washington School of Medicine, Seattle, WA, USA; Benaroya Research Institute, Seattle, WA, USA

**Keywords:** syphilis, secondary syphilis, CD4+ T cell, vaccine, tissue resident memory T cell, interferon gamma

## Abstract

**Background:** Histologic and serologic studies suggest the induction of local and systemic *Treponema pallidum* (*Tp*)-specific CD4+ T cell responses to *Tp* infection. We hypothesized that *Tp*-specific CD4+ T cells are detectable in blood and in the skin rash of secondary syphilis and persist in both compartments after treatment.

**Methods:** PBMC collected from 67 participants were screened by IFNγ ELISPOT response to *Tp* sonicate. *Tp*-reactive T cell lines from blood and skin were probed for responses to 88 recombinant *Tp* antigens. Peptide epitopes and HLA class II restriction were defined for selected antigens.

**Results:** We detected CD4+ T cell responses to *Tp* sonicate ex vivo. Using *Tp*-reactive T cell lines we observed recognition of 14 discrete proteins, 13 of which localize to bacterial membranes or the periplasmic space. After therapy, *Tp*-specific T cells persisted for at least 6 months in skin and 10 years in blood.

**Conclusions:** *Tp* infection elicits an antigen-specific CD4+ T cell response in blood and skin. *Tp*-specific CD4+ T cells persist as memory in both compartments long after curative therapy. The *Tp* antigenic targets we identified may be high priority vaccine candidates.

## INTRODUCTION

Despite widespread screening and treatment campaigns, the global incidence of syphilis has increased, with 7.1 million estimated new infections in 2020 [1]. In the United States, the rise is steepest among women, causing a doubling of cases of congenital syphilis since 2017 [2]. Experts concur that a vaccine against *Treponema pallidum (Tp)* would significantly reduce syphilis cases [3], but vaccine candidates have not yet advanced to clinical trials. Prototypes investigated in the rabbit model of experiment syphilis have focused on *Tp* outer membrane humoral targets and maximizing T_H_1 responses, and have elicited protection against lesion progression and dissemination [4].

It is hypothesized that during *Tp* infection, T_H_1-polarized CD4+ T cells produce interferon-γ (IFNγ) that activates macrophages to phagocytose and clear antibody-opsonized bacteria [5–9]. CD4+ T cells may also support treponemal clearance by providing B cell help to elicit opsonizing antibodies. Rabbit studies have yielded several T cell antigens [10–14], while fewer human data are available [15]. Abundant T_H_1-polarized CD4+ T cells are found in syphilitic lesions of humans and rabbits. However, antigens that stimulate T cells to produce IFNγ during human infection remain largely undefined.

In this study, we sought to identify *Tp* antigens recognized by CD4+ T cells during human infection. We hypothesized that B cell antigens and abundant *Tp* proteins would also contain T cell epitopes. Tissue resident-memory (T_RM_) are important for host protection against viral re-infection [16], but are less studied for bacterial infections. We further investigated the persistence of pathogen-specific T cells as skin T_RM_ after treatment and clinical resolution of secondary syphilis.

## METHODS

### Study Participants and Procedures

Participants recruited from the Public Health Seattle King County (PHSKC) Sexual Health Clinic (SHC) were enrolled at the University of Washington (UW) Virology Research Clinic. Criteria included clinical or serological evidence of syphilis, regardless of HIV infection status. Plasma HIV RNA, Rapid Plasma Reagin (RPR) and CD4+ T cell counts were measured in a clinical laboratory. Data were collected and managed using REDCap [17]. Studies were approved by the UW Institutional Review Board (STUDY00009493) and the PHSKC Research Advisory Council (20-689). Participants gave written informed consent.

We enrolled four cohorts: 1) persons with secondary syphilis (SS) defined as a compatible skin rash and positive RPR; 2) participants with asymptomatic (early and late latent) syphilis, defined as a positive treponemal and non-treponemal antibody results [18] but no clinical signs or symptoms of syphilis; 3) participants with a clinical history of previously-treated syphilis without clinical or serological evidence of new infection; and 4) participants with nonreactive treponemal and non-treponemal serology (no history of infection).

Blood was drawn within 72 hours of the clinic visit and provision of CDC guideline-compliant treatment [18]. Peripheral blood mononuclear cells (PBMC) were isolated and cryopreserved [19]. Three skin biopsies (3 mm) were collected from rash sites from participants (SS cohort) within 72 hours of treatment. Blood and biopsies of resolved rash site skin were also collected 30 and 180 days later.

### Antigens

#### Treponema pallidum

*Tp* sonicate was prepared from SS14 (Genbank NC_021508.1) treponemes expanded in co-culture with cottontail rabbit epithelial cells (Sf1Ep) for 8 days [20]. Sf1Ep cells with adherent and free treponemes were pooled after scrape-harvesting infected cultures. Trypsin was excluded to avoid degradation of *Tp* proteins. Treponemes were enumerated by dark field microscopy and bacteria pelleted at 23,000 x g for 30 minutes, 4°C. Mock-infected Sf1Ep cells were similarly harvested for use as control antigen. Pellets were resuspended to 1x10^9^ spirochetes/mL in sterile Dulbecco’s phosphate buffered saline (D-PBS) then sonicated at 10 kHz for 5 minutes on ice, cycling 10 seconds on/15 seconds off [21] with a 1/8 inch probe (QSonica N).

#### Proteins and peptides

The recombinant protein panel (n=88) included putative outer membrane proteins (OMP) [22,23], known targets of the humoral response in rabbits and humans [24–26], flagellar proteins, and proteins known to be abundant [27,28]. Full-length or truncated open reading frames (ORFs) were subcloned into pDEST203 **(Supplementary Tables 1 and 2)**. Selected additional *Tp* molecular clones (**Supplementary Table 3)** were included. A recombinant chimeric protein (p15/p17/p47) of Tp0171, Tp0435 and Tp0547, respectively, was also included (MyBioSource).

Synthetic peptides (13 amino acids overlapping by nine) were used for epitope mapping for Tp0435, Tp0548, Tp0574, Tp0684, Tp0856, Tp0858, and Tp0870 (70% purity, Genscript). Peptides were resuspended in dimethylsulfoxide (DMSO, Sigma) and studied as pools or single peptides with final peptide concentrations of 1 µg/mL maintaining a final DMSO concentration below 0.5% in all assays. Additional information on Methods is included in the **Supplementary Methods**.

### ELISPOT

Viable PBMC were counted (Guava, Luminex) after overnight rest for IFNγ ELISPOT (Mabtech) [19]. PBMC were plated at 2 x10^5^ cells/well in R10 medium [RPMI-1640, 10% fetal bovine serum (FBS), 2 mM L-glutamine, 100 U/mL penicillin/streptomycin (Gibco)]. *Tp* sonicate (1/10 dilution for a final concentration of 10^8^ *Tp*/mL), mock sonicate (diluted 1/10), 0.1 μg/mL p15/p17/p47 protein, R10 (negative control) and 1.6 μg/mL phytohemagglutinin (PHA-P, Remel) (positive control) were tested. Data acquired using an ImmunoSpot S6 Reader and ImmunoSpot Software (CTL) are reported as net spot forming units (SFU) per 10^6^ PBMC (average duplicate antigen response minus the average duplicate response to relevant negative control).

### Enrichment and expansion of *Tp*-specific T cells

*Tp*-reactive CD4+ T cells were enriched from PBMC using activation-induced marker (AIM)-based cell sorting, adapting published methods from other pathogens [19]. Cells were stained and live CD3+CD4+CD69+CD137+ cells were sorted and collected using a FACS Aria II (BD) **(Supplementary Figure 1A).**

T cells were expanded using standard methods [19,29]. Enrichment of *Tp*-specific T cells by AIM sorting was determined by comparing the net (*Tp* minus mock) CD4+ T cell responses ex vivo during AIM to the net *Tp*-reactive cells determined by intracellular cytokine (IFNγ and IL-2) staining **(Supplementary Table 4 and Supplemental Figure 1).**

### T cell antigen specificity assays

TCL were screened in duplicate. Equal numbers of responder T cells (10^5^) and antigen presenting cells (APC) were combined with antigens in 200 μL TCM. γ-irradiated autologous PBMC were used as APC for IVTT (in vitro transcription/translation)-generated proteins (1/2,000 final dilution). Epstein-Barr virus-transformed lymphoblastoid cell lines (LCL) prepared from PBMC [19] were used as APC for synthetic peptides (1 μg/mL). *Tp (*5x10^7^ *Tp*/mL) and mock sonicate (1/20 dilution) were added as positive and negative controls, respectively. At three days, [^3^H] thymidine (1 μCi/well) was added; cells were harvested after 18h and counted by scintillation (TopCount, Perkin Elmer) [16]. TCL were considered reactive for *Tp* antigens if the proliferative response for both technical replicates (counts per minute, CPM) exceeded a statistically determined threshold (below) in two independent assays.

### HLA restriction

HLA typing was performed for selected participants (Scisco) [30]. HLA restricting loci were determined using peptide titration (0.01, 0.1, and 1 μg/mL peptide) in the presence of blocking anti-HLA-DR, -DP, or -DQ antibodies [29] and autologous LCL (5x10^4^ cells per well). To identify HLA restricting alleles, artificial APC (aAPC) (3x10^4^) expressing single participant-relevant HLA class II (or no HLA) were co-cultured with T cells (10^5^) and peptides (1 μg/mL) [31]. After 24 hours, supernatant IFNγ was measured by ELISA [32].

### Statistics

The threshold for positivity in proliferation assays was calculated as (median response of all negative control antigens) + [2.33 times median absolute deviation (MAD) of all negative control antigens] for a false discovery rate <1% [33,34]. Kruskal-Wallis with Dunn’s post hoc test for multiple comparisons was used to compare groups for ELISPOT data. Linear and logistic regression was used for multivariable analyses. Two-sided P values <0.05 were considered statistically significant. Statistical tests were performed using Prism (v10, GraphPad).

## RESULTS

### Participants

The 67 participants **(Table 1)** included persons with clinically and/or serologically diagnosed syphilis or control individuals. Demographic characteristics reflect persons visiting the SHC (79% male, 44% White) [35]. Women represented 66.7% of the control group with no history of infection and 20.6% of the syphilis group. The median RPR titer was lower in the treated than in the secondary syphilis group (1:4 vs 1:128). Participants with multiple prior syphilis infections were common, especially among asymptomatic seropositive participants. Participant 17 was initially enrolled with secondary syphilis, treated with 4-fold RPR titer decline and resolution of symptoms at 180 days, and re-enrolled with early latent syphilis 10 months later (day 480). HIV co-infection was present in eight participants (16%) with prior or current *Tp* infection. All participants with HIV infection were virologically suppressed on antiretroviral therapy, with blood CD4+ T cell counts >300 per μL.

**Table 1.**
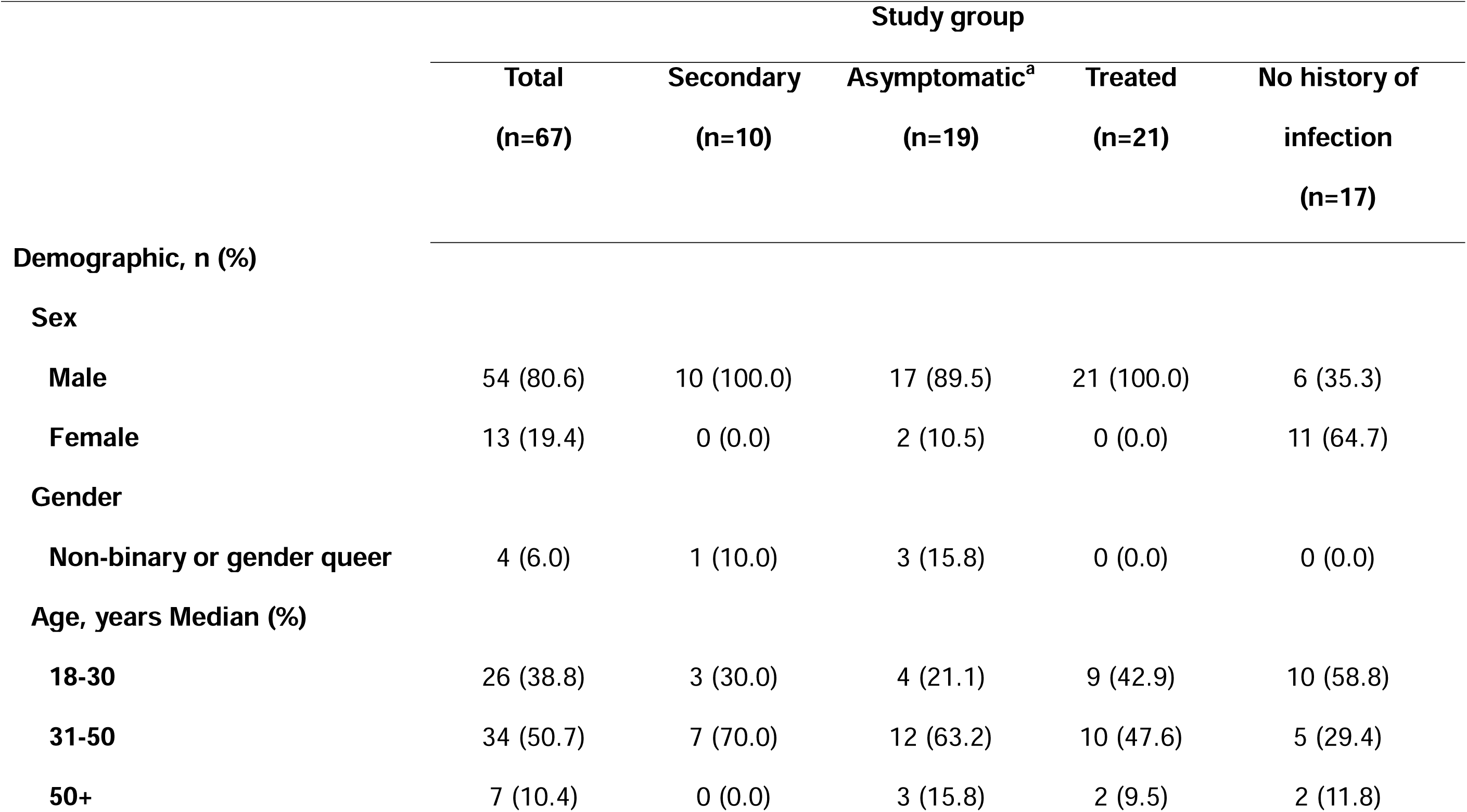
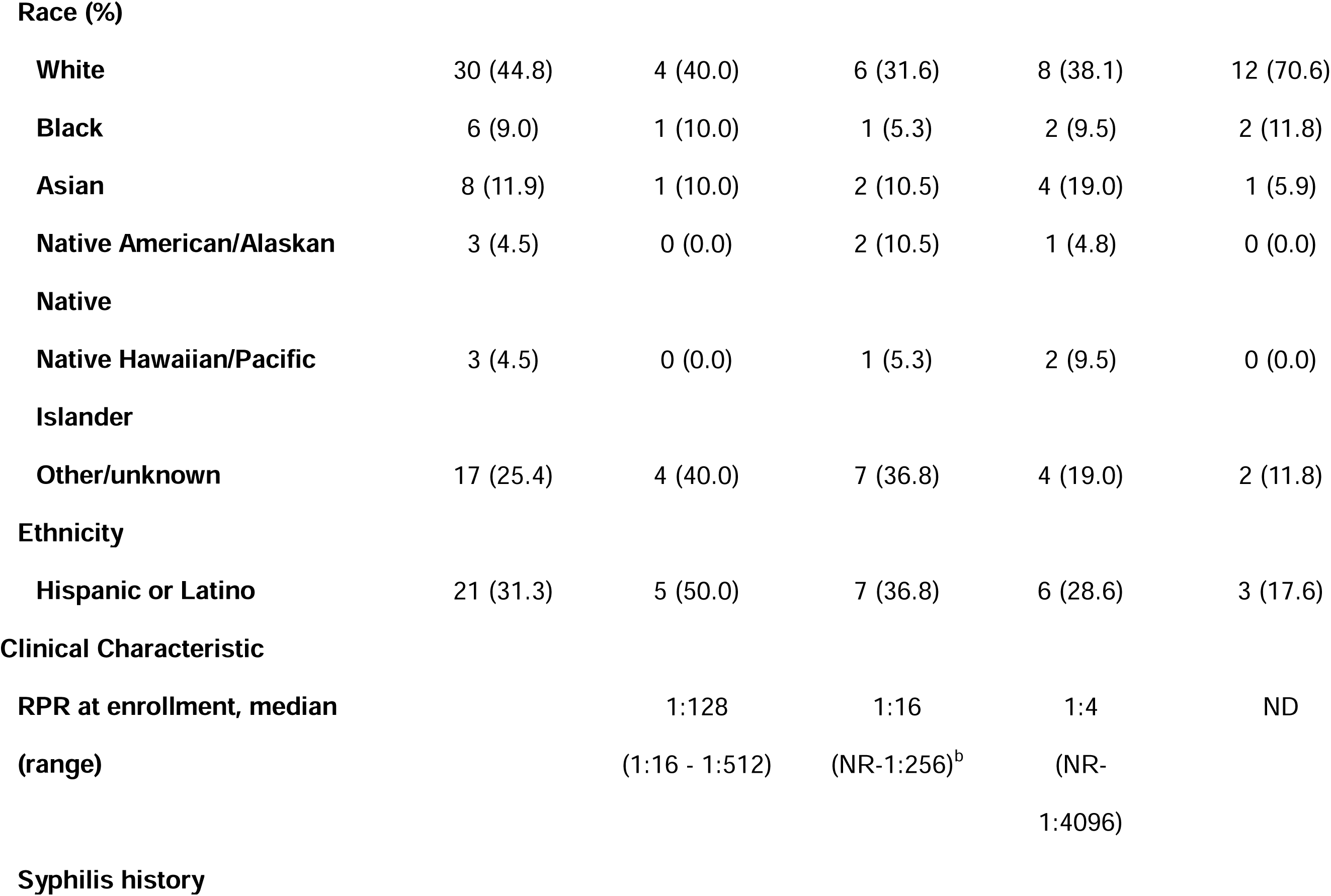
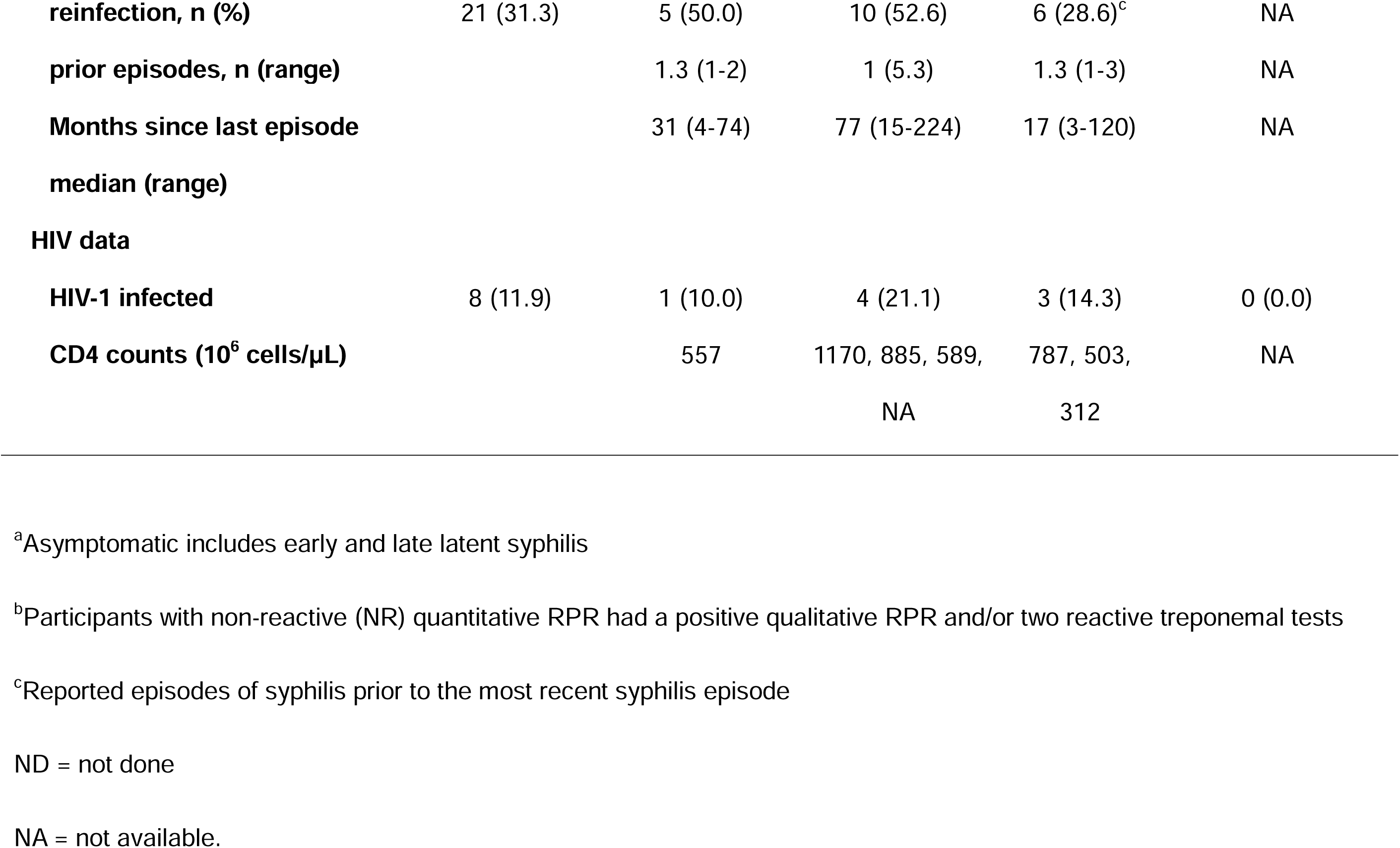
Clinical and demographic characteristics of study participants.

### PBMC produce IFNγ in response to *Tp* sonicate

Using ex vivo participant PBMC, we evaluated cellular responses to *Tp* sonicate by IFNγ ELISPOT. The median response in the previously treated group was significantly higher (37 SFU/10^6^ PBMC, interquartile range (IQR) 17-123) than the group with no history of infection (6 SFU/10^6^ PBMC, IQR 0.1-24) (p=0.003). Interestingly, participants with secondary and latent syphilis did not have higher responses than participants with no history of infection **(Figure 1A).** Notably, HIV infection did not prevent detection of PBMC responses to *Tp* infection. Responses in the no history of infection group were generally less than 15 SFU/10^6^ PBMC.

**Figure 1.**
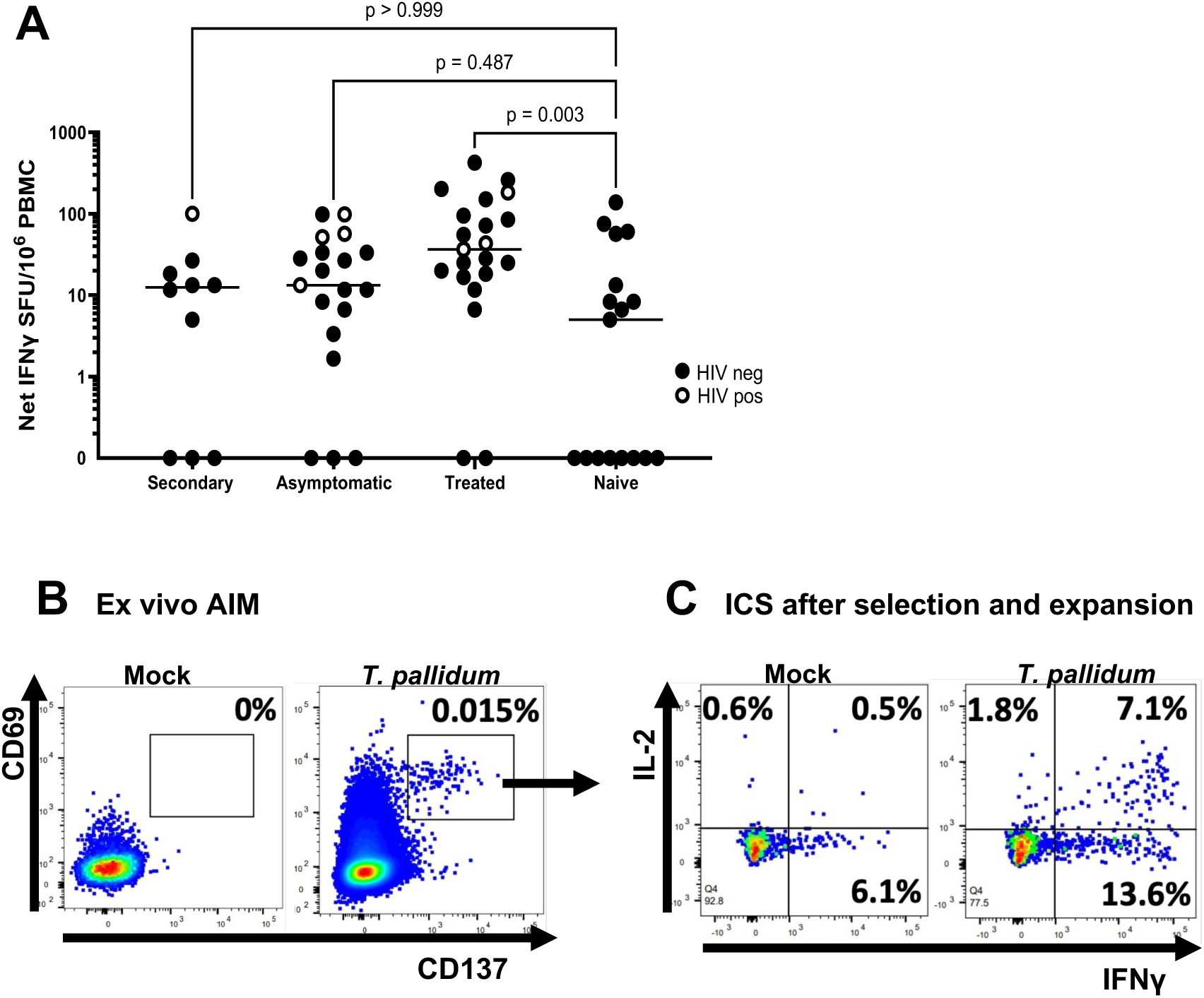
*A,* T cell response to *Tp* sonicate. PBMC were incubated with *Tp* or mock sonicate. Activation was measured by IFNγ ELISPOT. Data points are the average of duplicates. White dots represent HIV-infected participants. Horizontal lines are median SFU/million PBMC. p values were calculated by Kruskal-Wallis test between groups. *B, C,* Enrichment of rare *Tp*-reactive CD4+ T cells from PBMC. *B,* PBMC from participant 02, 47 months after treatment for early latent syphilis were stimulated with the indicated antigens. Gated live CD3+CD4+CD8- T cells were analyzed for activation. Cells in the indicated gate were sorted and bulk-expanded. Percentages of activated cells are indicated. *C,* The resultant polyclonal CD4+ T cell line was tested for reactivity to *Tp* or mock sonicate. Dot plots show gated live CD3+CD4+CD8- responder cells. Percentages of cytokine positive responder cells are indicated.

After infection, individuals develop a lifelong antibody response to specific *Tp* proteins used in serology assays. Reflecting the cognate CD4+ T-B cell theory of T cell help, we expected T cell responses to a p15/p17/p47 fusion protein used for serology. We observed such responses, and a positive correlation to whole *Tp* sonicate (R 0.53, n=60, p<0.001) **(Supplementary Figure 2)**.

### Rare circulating CD4+ T cell respond to *Tp* sonicate

PBMC specimens with a positive ELISPOT response to *Tp* were prioritized for further study. Since diverse populations of T and NK cells can express IFNγ, we used activation-induced markers (AIM) expression to focus specifically on CD4+ T cells **Figure 1B)**. Amongst ex vivo PBMC, a median of 0.02% (IQR 0.003-0.18%) of CD4+ T cells were *Tp*-reactive **(Table 2)**.

**Table 2.** Abundance of *Tp*-reactive CD4+ T cells in PBMC and after enrichment.

| **Participant ID** | **Cohort** | ***Tp*-specific CD4+ T cells, (%) ex vivo<sup>a</sup>** | ***Tp*-specific CD4+ T cells, (%) after enrichment<sup>b</sup>** | **Fold enrichment<sup>c</sup>** |
| --- | --- | --- | --- | --- |
| **13** | Secondary | 0.010 | 1.09 | 109 |
| **14D0** | Secondary | 0.023 | 9.10 | 401 |
| **14D30<sup>d</sup>** | Secondary | 0.029 | 3.70 | 126 |
| **14D180<sup>d</sup>** | Secondary | 0.074 | 4.40 | 59 |
| **15** | Secondary | 0.020 | 0 | NA |
| **16** | Secondary | 0.120 | 0 | NA |
| **17D0** | Secondary | 0.070 | 0 | NA |
| **17D30<sup>d</sup>** | Secondary | 0.113 | 2.40 | 21 |
| **17D180<sup>d</sup>** | Secondary | 0.039 | 14.30 | 369 |
| **17D480<sup>e</sup>** | Asymptomatic | 0.010 | 36.30 | 3630 |
| **18D0** | Secondary | 0.006 | 0 | NA |
| **18D30<sup>d</sup>** | Secondary | 0.006 | 0 | NA |
| **10** | Asymptomatic | 0.011 | 0 | NA |
| **12** | Asymptomatic | 0.009 | 22.00 | 2444 |
| **19** | Asymptomatic | 0.038 | 0 | NA |
| **01** | Treated | 0.036 | 0 | NA |
| **02** | Treated | 0.015 | 15.30 | 1020 |
| **03** | Treated | 0.012 | 0.20 | 17 |
| **04** | Treated | 0.005 | 4.20 | 784 |
| **05** | Treated | 0.003 | 3.70 | 1088 |
| **06** | Treated | 0.176 | 3.00 | 17 |
| **07** | Treated | 0.132 | 23.50 | 178 |
| **08** | Treated | 0.019 | 32.90 | 1759 |
| **09** | Treated | 0.020 | 2.83 | 142 |
| **09D210<sup>f</sup>** | Asymptomatic | 0.030 | 3.84 | 128 |
| **11** | Treated | 0.088 | 0.50 | 6 |

<sup>a</sup>Net percent of gated live CD4+ T cells expressing CD69 and CD137 after stimulation with *Tp* sonicate
<sup>b</sup>Net percent of gated live CD4+ responder T cells expressing IFNγ , IL-2 or both cytokines after stimulation with *Tp* sonicate
<sup>c</sup>Fold enrichment of *Tp*-specific CD4+ T cells
<sup>d</sup>PBMC collected the indicated day after treatment
<sup>e</sup>RPR rise was diagnostic for re-infection between days 180 and 480
<sup>f</sup>RPR rise was diagnostic for re-infection between initial enrollment and day 210
NA = not applicable.

### *Tp* proteins recognized by circulating and skin resident CD4+ T cells

To enable testing of large antigen sets, live antigen-reactive CD4+ T cells were functionally enriched by sorting AIM+ cells and polyclonal expansion. As measured by ICS, *Tp*-specific CD4+ T cells were enriched ∼1000-fold in an illustrative participant **(Figures 1B and 1C**). Amongst our cohort of *Tp*-infected persons overall, the median reactivity of enriched TCL to *Tp* sonicate was 2.9% (IQR 0.0-36.3%) of CD4+ T cells (**Table 2).** TCL from eight participants with diverse clinical status were selected for further study based on reactivity to *Tp* sonicate and the availability of multiple time points. We screened TCL with 88 recombinant *Tp* proteins **(Supplementary Tables 1-3)** in a proliferation assay. Proteins that elicited responses in at least two independent assays were reported as positive. Reactivity to one or more recombinant *Tp* antigens was detected in five of eight participants studied [example of raw data showing reactivity to Tp0621, Tp0769, and Tp0792 **(Figure 2A)**]. For comparisons across multiple TCL, we normalized the results for individual TCL to the median response to negative control antigens **(Statistics; representative data, Figure 2B and Supplementary Table 4)**.

**Figure 2.**
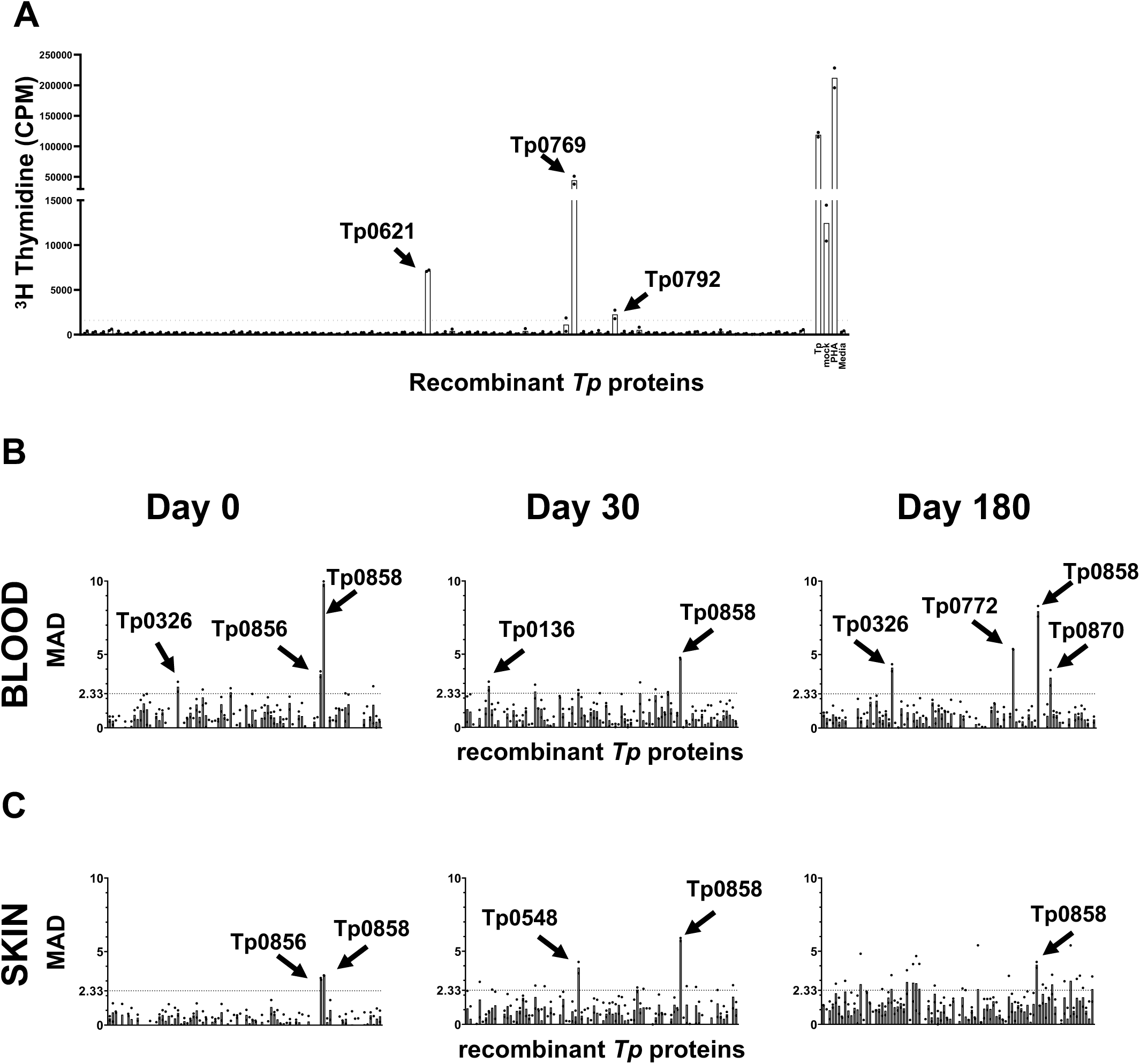
Identification of *Tp* CD4+ T cell antigens. TCL were enriched and expanded from blood (*A, B*) or cultured directly from skin (*C*) on day 0, day 30 and day 180 after treatment. *A*, TCL proliferation in response to defined *Tp* antigens is reported as counts per minute from participant 17 with early latent syphilis. Controls at right include *Tp* sonicate (*Tp*), mock sonicate (mock) and PHA. *B and C,* Serial T cell responses to *Tp* proteins in blood and skin from participant 14 with secondary syphilis. Data are reported as median absolute deviation (MAD) above the median proliferative response to unrelated recombinant proteins. The positivity threshold is indicated by dashed horizontal lines. Technical duplicates are shown as black dots and bars are means. Responses are designated positive if both replicates are above threshold. *Tp* antigens with positive responses are labeled.

The presence and specificity of *Tp*-specific CD4+ T cells in skin was studied in two participants (14 and 17) at the diagnosis of secondary syphilis and 30 and 180 days after treatment. For these subjects, skin lesion-derived T cells were also polyclonally expanded, without preliminary in vitro enrichment, based on the hypothesis that bacteria-reactive lymphocytes might be naturally localized to infected areas. *Tp*-specific T cells were readily detected by proliferation assay in skin-derived TCL. For participant 14, we observed agreement in *Tp* antigens recognized between skin and PBMC-derived TCL. We also detected consistency in the *Tp* antigens driving CD4+ T cell responses over time **(Figures 2B and 2C)**. Durable antigens included the predicted OMP FadL family proteins Tp0548, Tp0856 and Tp0858 [22]. In contrast, subject 17 showed differences over time. We observed responses to *Tp* sonicate during initial infection, but PBMC-derived TCL responses to members of our *Tp* protein panel were only observed during an asymptomatic reinfection that occurred 10 months later, at which time Tp0621, Tp0769, and Tp0792 were reactive. **(Figures 2A and 3A)**. Integrating cross-sectional and longitudinal blood and skin specimens, we detected CD4+ T cell responses to 14 distinct *Tp* proteins (**Figure 3A and Supplementary Table 5**). Four of the 14 proteins were recognized by multiple TCL. Many are predicted to reside in the inner or outer cell membrane or periplasmic space **(Figure 3B).**

**Figure 3.**
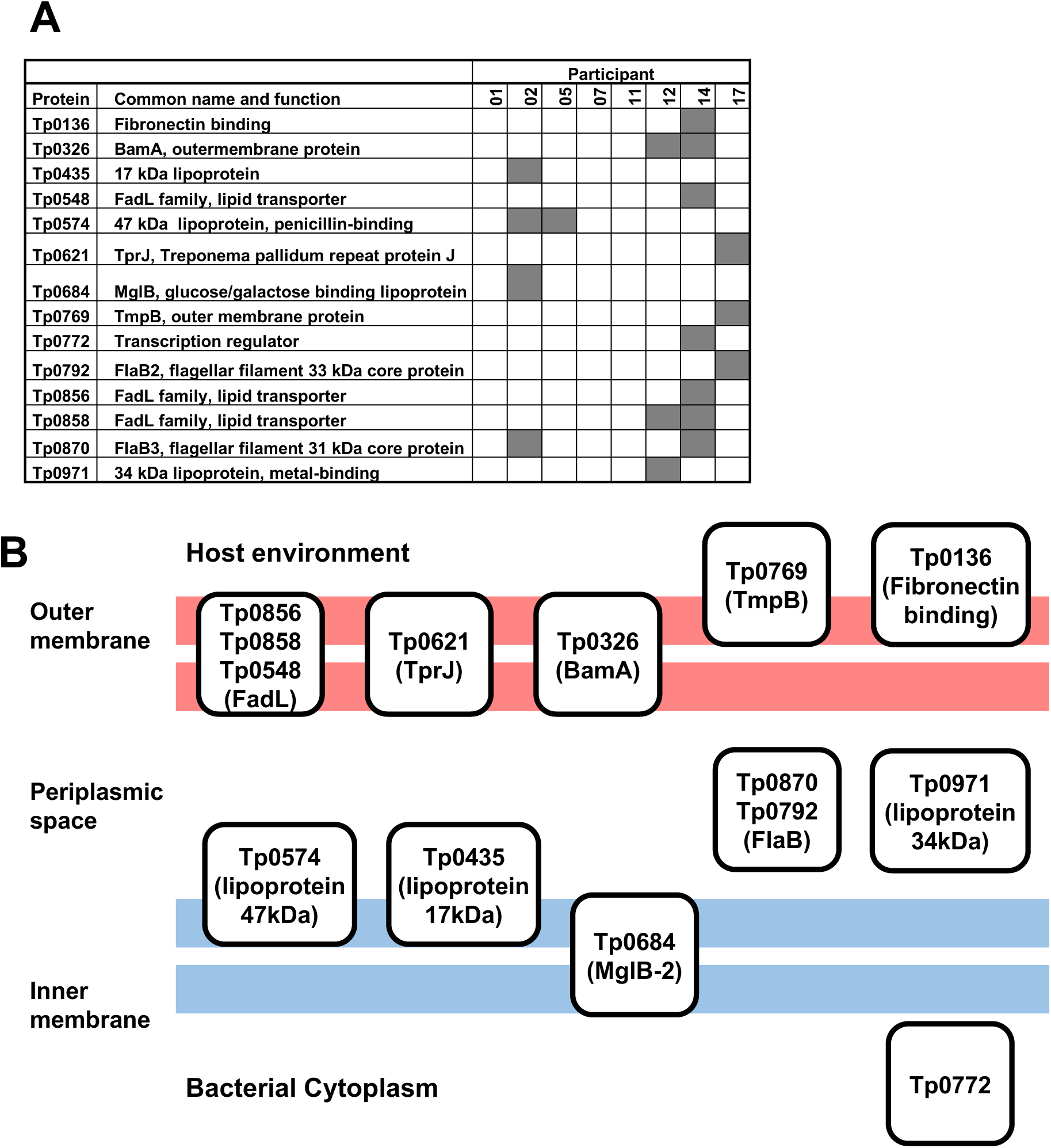
Summary of CD4+ T cell antigen discovery. *A, Tp* proteins driving T cell responses. For participants with more than one PBMC- or skin-derived TCL tested, results are summarized per participant. *B,* Predicted localization of *Tp* CD4+ T cell antigens in the bacterial cell.

### Presentation of *Tp* epitopes involves HLA-DR, -DP, and -DQ loci

To confirm that the reactivity we observed to *Tp* proteins was classically peptide- and HLA-restricted, we performed limited epitope mapping. PBMC-derived CD4+ TCL or skin TCL served as responder cells, and autologous LCL as antigen presenting cells. We tested peptide pools as and then deconvoluted active pools to individual peptides **(Supplementary Figures 3A, 3B, 4 and 5)**. Responses were detected (representative data, **Figure 4A and Supplementary Figure 3)** to 16 unique CD4+ T cell epitopes in seven *Tp* proteins from three participants (**Figure 4D)**. We determined the HLA class II restriction for selected epitopes. First, using locus-specific HLA class II blocking mAbs, we confirmed that nine epitopes were restricted by HLA-DR, two were restricted by HLA-DP, and one by HLA-DQ **(Supplementary Figure 3C, Figure 4D)**. For example, responses to a peptide in Tp0684 **(Figure 4A)** were blocked by anti-HLA-DR **(Figure 4B)**, but not by anti-HLA-DP or -DQ. The Tp0684 peptide was further tested using aAPC expressing a single participant-relevant HLA. TCL were activated in the presence of HLA-DRB1*07:01, but not HLA-DRB1*15:01, HLA-DRB5*01:01, or negative control cells **(Figure 4C)**. Overall, CD4+ T cell reactivity to the *Tp* antigens studied involves classical antigen presentation by HLA class II.

**Figure 4.**
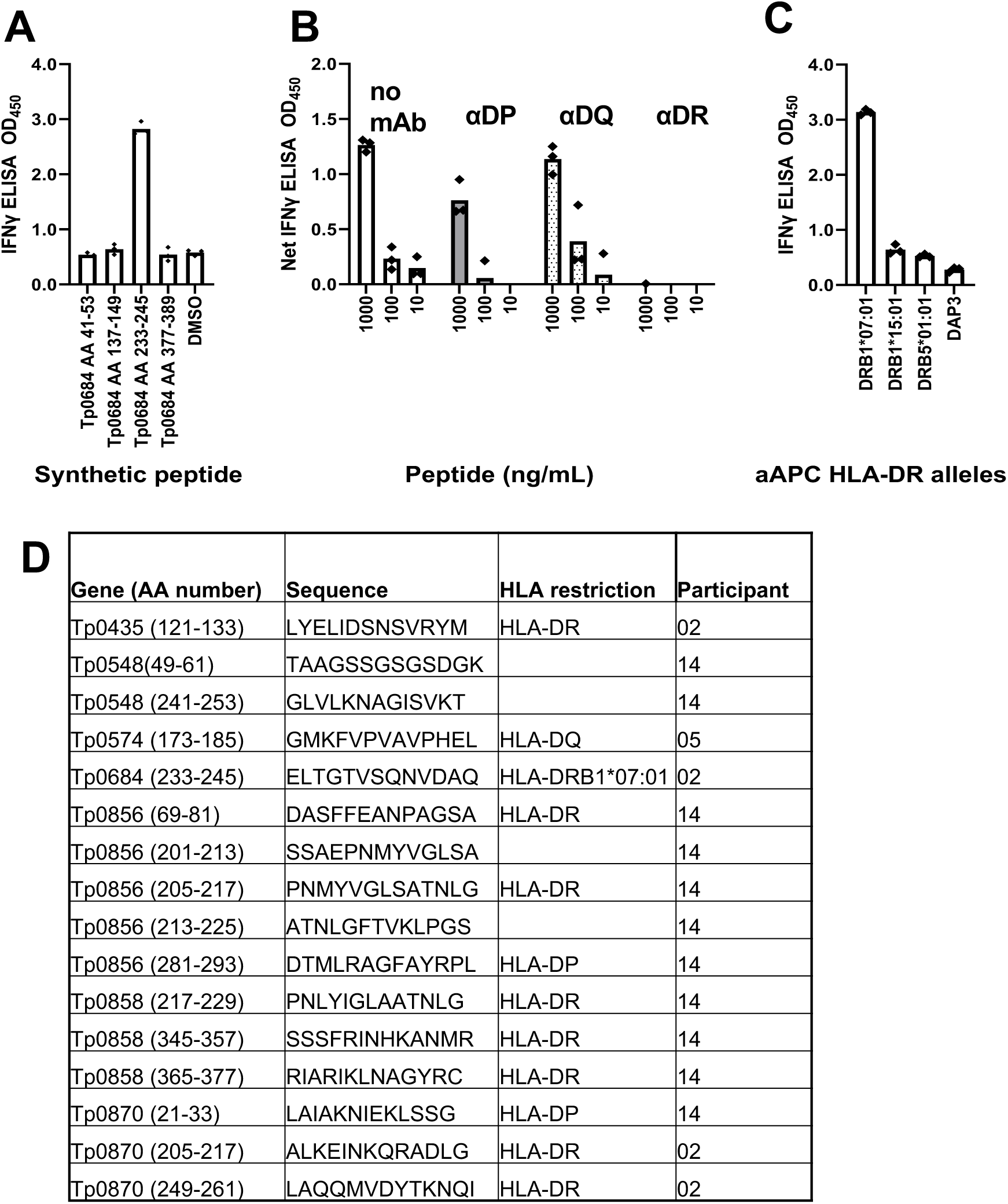
CD4+ T cell epitope mapping and determination of HLA restriction. Representative data from participant 02 who was treated for early latent syphilis 47 months before sampling. *A,* PBMC-derived TCL reactivity to Tp0684 (MglB-2) peptides at 1 μg/mL. Triplicate raw data for the active peptide and representative non-reactive peptides are shown. *B,* Reactivity of polyclonal *Tp* -reactive CD4+ T cells to peptide Tp0684 AA 233-245C. Peptide is titrated from 1 to 0.01 μg/ml. HLA class II restriction is determined by mAb blocking with mAb specificity shown at top. *C,* IFNγ secretion by the same effector cells in response to 1 μg/mL peptide Tp0684 AA 233-245 using aAPC expressing participant-relevant HLA DR alleles and the control parent cell line, DAP3. *D,* Summary of T cell epitopes and available HLA restriction data.

## DISCUSSION

Little is understood about the T cell response to *T. pallidum* during human infection. Syphilis has distinct temporal phases from primary to tertiary infections with specific diagnostics and therapeutics. Because secondary syphilis presents several weeks after infection and induction of acquired immunity, and because secondary syphilis skin lesions resolve without antibiotics, we hypothesized that circulating *Tp*-specific T cells would be detectable when secondary syphilis presents, and that local *Tp*-reactive T cells might be present and possibly contributing to bacterial clearance. Our observations are consistent with these hypotheses and with earlier findings that CD4+ T cell infiltrates are readily detectable in secondary syphilis lesions [8] and that T_H_1 cytokines are produced locally in lesions [6]. Our study also sheds light on immune memory to syphilis. We found that *Tp*-specific CD4+ T cells, although rare amongst PBMC, continue to respond to defined *Tp* antigens years after syphilis treatment, consistent with development of a durable memory response. Interestingly, PBMC IFNγ responses to *Tp* during secondary and asymptomatic syphilis were not significantly different than the group with no history of infection. This may be explained by the observation that pathogen-specific T cells are sequestered at sites of infection and not circulating [21,36]. Indeed, we found that effector T_H_1 CD4+ T cells were enriched in active secondary syphilis skin biopsies and readily detectable without an in vitro enrichment step. We also observed *Tp*-reactive CD4+ T cells for at least six months after antibiotic therapy, suggesting local retention as T_RM_.

*Tp* infects primarily through mucosa and breaks in epithelia. Paired collection of skin biopsies and PBMC allowed us to compare the circulating and local T cell responses to *Tp* infection during active infection and months after treatment. We found congruence between PBMC and skin T cell antigenic specificity, suggesting seeding of T_RM_ by circulating cells that recognize outer membrane antigens. Elicitation of T_RM_ at skin and mucosal sites through immunization may enhance protection against tissue invasion, clinical illness, and transmission.

In the present study, we define the fine specificity of *Tp*-reactive CD4+ T cells. Based on cognate T cell help theory, we reasoned that CD4+ T cells may recognize known B cell antigens [37]. We identified the diagnostic antigens Tp0435 and Tp0547, but not Tp0171, as stimulators of long-lived (>10 years after treatment) CD4+ T cell responses. Our data may also identify or prioritize potential vaccine antigens. Periplasmic FlaB3 (Tp0870) [38,39] and the lactoferrin-binding protein,Tp34 (Tp0971) [40], were identified as a human CD4+ T cell antigens. When used as immunogens, these proteins induce partial protection in rabbits by humoral and cellular immune responses. The glucose transporter, MglB-2 (Tp0684), is another known antibody target [41,24] that was identified as a T cell antigen.

In addition to known antibody targets, we focused on other OMP, with the rationale being their potential to elicit opsonic antibodies. Despite the outer membrane of *Tp* having relatively few proteins [42], we identified multiple predicted OMP as T cell antigens. These include the fibronectin binding protein, Tp0136 which was recently shown to alter clinical progression in the rabbit model after immunization and is T cell antigenic in this species [43,44]. Of the *Tp* repeat (Tpr) family, which includes targets of opsonophagocytosis,the TprJ (Tp0621) fragment [14] was identified as a T cell antigen. Other B cell antigens that may facilitate opsonophagocytosis and stimulate human T cells include the OMP Tp0326 [45], Tp0769, and members of the FadL lipid transporter family. For example, Tp0858 is a FadL protein and is highly expressed in *T. pallidum* [27,28] and has immunogenic extracellular loops [46]. We observed that T cell responses to this protein persisted in blood and skin. In addition, our analyses reveal partial overlap between T cell antigens reported in rabbits and those we report in humans, including Tp0136, Tp0621, Tp0870, Tp0574, and Tp0435 [10,11,14,39,44]. These data bridge the gap between the experimental animal model for *Tp* infection and the natural human host.

We detected blood *Tp*-specific CD4+ T cell responses at syphilis diagnosis and long after treatment. Syphilis reinfection is known to occur, such that memory immune responses are not entirely protective against challenge. For example, participant 14 was enrolled after multiple episodes of prior symptomatic syphilis. Intriguingly, the emergence of detectable antigen-specific T cell responses in participant 17 during an asymptomatic reinfection suggests a relationship between memory *Tp*-specific T cells and reduced disease burden. The concept of partial protection is supported by the clinical observations that reinfections are more likely than primary infections to be asymptomatic [47]. Further research is required to understand the kinetics of systemic and local (skin) T cell responses and the immune correlates of infection severity.

Our study had strengths and limitations. Paired, longitudinal blood and skin samples allowed us to document that *Tp*-specific T cells participate in the acute inflammatory infiltrate in secondary syphilis and persist locally as T_RM_. Despite the low abundance of *Tp*-reactive T cells in PBMC, we were able to query responses at the antigen- and peptide-specific levels using AIM-based enrichment. The use of in vitro T cell polyclonal expansion introduces the potential for biased clonal expansion, and further limits measurement of specific antigens to a qualitative yes/no call rather than a quantitative level. The assays employed in this study were limited to detection of T_H_1 CD4+ T cell responses, but other CD4+ T cell subsets and perhaps CD8+ T cells may also have protective roles [8]. Our small sample size is another limitation and precludes us from generalizing about the CD4+ T cell epitopes in larger populations. The HLA-DRB1*07:01 allele is prevalent globally, suggesting that additional epitope discovery could facilitate the design of vaccine antigens with wide CD4+ T cell population coverage [48].

Our protein antigens were designed from the Nichols strain and covered only 8% of the genome (88 of 1039 predicted open reading frames). Conversely, half of the genomes isolated from participants in our geographic location are in the SS14 clade of treponemes [49], which we used for initial enrichment of T cells from blood. Although the Nichols and SS14 genomes are >99% identical [50] and strain-specific T cell epitopes are unlikely [12], we could have missed such responses using Nichols-derived proteins. Comparisons of our identified T cell epitopes amongst 351 annotated *Tp* genomes disclosed only two protein coding variants: in amino acid 241-253 of Tp0548, which is known to have inter-strain variability [50], and in the protein Tp0435. While antigenic variation is unlikely to contribute to the response patterns we noted, decreased sensitivity to detect T cell activity could result from shallow sampling of the *Tp*-reactive CD4+ T cell repertoire related to the initial blood volume, utilizing only CD69 and CD137 as AIMs, and variable proliferation of *Tp*-reactive T cells in vitro. Future work will seek to rectify these factors by starting with increased numbers of AIM+ cells selected with an expanded panel of activation markers, and ultimately progression to direct ex vivo assays.

In summary, we show that *Tp* infection elicits an antigen-specific CD4+ T cell response in both blood and skin and that *Tp*-specific T cells can be retained in both compartments long after curative therapy. These findings suggest the importance combining data from *Tp*-infected humans and rabbits to inform the identification of vaccine candidates to be prioritized for clinical studies. Further work is needed to expand the known repertoire of human CD4+ T cell epitopes recognized during syphilis infection and to characterize T cell responses that protect against *Tp* reinfection.

## Acknowledgments

The authors thank the study participants and clinicians and Dr. Sheila Lukehart for stimulating discussions and manuscript review.

## Data available in supplementary material

### Foot note page

Potential conflicts of interest (COI) are listed. D.M.K. reports membership on the Scientific Advisory Board of MaxHealth LLC and Curevo Vaccines, grant support from Sanofi Pasteur, and co-inventorship of institutionally owned patents concerning HSV vaccines. All authors report no potential conflicts and have submitted the ICMJE Form for Disclosure of Potential Conflicts of Interest.

Financial support. This work was supported by the National Institute of Allergy and Infectious Diseases, National Institutes of Health (U19AI144133 to A.W., C.E.C., L.G., A.L.G., T.B.R. and D.M.K. T.B.R. was also supported on UM1AI148684 and T32AI07140, K.L.H. was supported on U19AI144177). The content is solely the responsibility of the authors and does not represent the official views of the National Institutes of Health. Redcap software is supported by NIH award UL1TR002319.

