## Supplementary Materials for "*Treponema pallidum* periplasmic and membrane proteins are recognized by circulating and skin CD4+ T cells"

Supplementary Tables.

Supplementary Table 1. PCR primers for amplification and cloning into pDEST203

| *Tp* locus | other name(s) | 5' Primer (5' to 3')^a^ | 3' Primer (5' to 3')^a^ | NCBI ID | Minimum | Maximum | Length (nt) |
| --- | --- | --- | --- | --- | --- | --- | --- |
| Tp0017 | tetratricopeptide repeat protein CDS | ggggacaagtttgtacaaaaaagcaggcttcATGGTTGCACGATCGATTCGT | ggggaccactttgtacaagaaagctgggtcCGCTATCCTCACATAGCGAT | AE000520.1 | 18404 | 19348 | 945 |
| Tp0126 | hypothetical protein CDS | ggggacaagtttgtacaaaaaagcaggcttcATGCGGACGCACGACATACCCCGC | ggggaccactttgtacaagaaagctgggtcGAGGTGGTAGCGAACGTCCAC | WP_014342713.1 | 147623 | 148294 | 672 |
| Tp0131 | Tpr protein D CDS | ggggacaagtttgtacaaaaaagcaggcttcGTGGGCAGGCAGGTG | ggggaccactttgtacaagaaagctgggtcTTACCATGTCACTTTCATTCCG | WP_010881566.1 | 134912 | 136708 | 1797 |
| Tp0155 | M23 family metallopeptidase CDS | ggggacaagtttgtacaaaaaagcaggcttcGTGAGCGTGTACTGTCGAAGTTC | ggggaccactttgtacaagaaagctgggtcCGGAAGGGTACGCATACGCAG | WP_010881602.1 | 178280 | 179395 | 1116 |
| Tp0171 | FMN-binding protein CDS | ggggacaagtttgtacaaaaaagcaggcttcATGGTGAAAAGAGGTGGC | ggggaccactttgtacaagaaagctgggtcCTACCTGCTAATAATGGCTTC | WP_010881618.1 | 192146 | 192574 | 429 |
| Tp0249 | flagellar filament outer layer protein CDS | ggggacaagtttgtacaaaaaagcaggcttcATGAAGAAAGCGGTTGTGTTG | ggggaccactttgtacaagaaagctgggtcCTACTGCTGCTCTTCCTGC | WP_010881697.1 | 262600 | 263652 | 1053 |
| Tp0257 | glpQ CDS, protein D, glycerophosphodiester phosphodiesterase | ggggacaagtttgtacaaaaaagcaggcttcATGCGGGGAACATATTGTG | ggggaccactttgtacaagaaagctgggtcTCAATAGCGGGCGGG | WP_010881706.1 | 269345 | 270415 | 1071 |
| Tp0277 | S41 family peptidase CDS | ggggacaagtttgtacaaaaaagcaggcttcATGCAGACGGTGCAGG | ggggaccactttgtacaagaaagctgggtcTCAAGATACCTTCTTTTTTTCCTG | WP_010881726.1 | 293751 | 295097 | 1347 |
| Tp0315 | DUF2715 domain-containing protein CDS | ggggacaagtttgtacaaaaaagcaggcttcTTGGTGAGGGGTTGTCGTGTG | ggggaccactttgtacaagaaagctgggtcATACGCCGACTCTTCGTG | [WP_010881764.1](https://www-ncbi-nlm-nih-gov.offcampus.lib.washington.edu/protein/499184224) | 331661 | 332306 | 645 |
| Tp0398 | fliE CDS | ggggacaagtttgtacaaaaaagcaggcttcATGACGCCAGTTGGTACC | ggggaccactttgtacaagaaagctgggtcTTACCGAGCAGTGGTGAC | WP_010881846.1 | 423933 | 424307 | 375 |
| Tp0400 | fliG CDS | ggggacaagtttgtacaaaaaagcaggcttcATGGCCGTTACATCCGTG | ggggaccactttgtacaagaaagctgggtcTTACACAATCATCTCGTC | WP_010881848.1 | 426132 | 427190 | 1059 |
| Tp0435 | copper resistance protein NlpE CDS | ggggacaagtttgtacaaaaaagcaggcttcATGAAAGGATCTGTCCGCG | ggggaccactttgtacaagaaagctgggtcCTATTTCTTTGTTTTTTTGAGCAC | WP_010881883.1 | 463606 | 464076 | 471 |
| Tp0453 | membrane protein CDS | ggggacaagtttgtacaaaaaagcaggcttcTTGATAAGGCGTAGATATCGTGG | ggggaccactttgtacaagaaagctgggtcTTACGAACTTCCCTTTTTGGAG | WP_010881902.1 | 483698 | 484561 | 864 |
| Tp0483 | fibronectin type III domain-containing protein CDS | ggggacaagtttgtacaaaaaagcaggcttcATGAGTATCATTAGCCGCGTG | ggggaccactttgtacaagaaagctgggtcGTTATGAAAGCGATAGCCGAACG | WP_014342802.1 | 514108 | 515319 | 1212 |
| Tp0548 | UPF0164 family protein CDS | ggggacaagtttgtacaaaaaagcaggcttcATGCGTCAGAATGGC | ggggaccactttgtacaagaaagctgggtcCTAGTTGAGAATATCAAAGAGAG | WP_010881995.1 | 593270 | 594574 | 1305 |
| Tp0574 | hypothetical protein CDS | ggggacaagtttgtacaaaaaagcaggcttcGTGAAAGTGAAATACGCAC | ggggaccactttgtacaagaaagctgggtcCTGGGCCACTACCTTCGCACGC | WP_010882021.1 | 623858 | 625162 | 1305 |
| Tp0610 | hypothetical protein CDS, TprH | ggggacaagtttgtacaaaaaagcaggcttcATGAAAGGAGTACGCTGTCC | ggggaccactttgtacaagaaagctgggtcCCAGTTAATAGTCATTCCGC | WP_010882056.1 | 662845 | 664926 | 2082 |
| Tp0663 | flagellar filament outer layer protein FlaA CDS, Tromp2 | ggggacaagtttgtacaaaaaagcaggcttcATGAAACAGGGCTG | ggggaccactttgtacaagaaagctgggtcTCATTTGCCGCTCTCTCC | WP_014342815.1 | 727147 | 727875 | 729 |
| Tp0684 | substrate-binding domain-containing protein CDS, MglB-2 | ggggacaagtttgtacaaaaaagcaggcttcATGAAGGAGAATTCTTGCACG | ggggaccactttgtacaagaaagctgggtcTCAGTATTTGAGCTTGTCTG | WP_010882129.1 | 749861 | 751072 | 1212 |
| Tp0698 | DUF2715 domain-containing protein CDS | ggggacaagtttgtacaaaaaagcaggcttcATGAGAAGATTGCTGGCATG | ggggaccactttgtacaagaaagctgggtcCACTCGGAACGTCGGCCC | WP_010882143.1 | 767377 | 767922 | 546 |
| Tp0727 | flgE CDS | ggggacaagtttgtacaaaaaagcaggcttcATGATGCGGTCACTTTTTTCAG | ggggaccactttgtacaagaaagctgggtcTCAGCGCTTCAAACTCAAC | WP_010882172.1 | 793546 | 794937 | 1392 |
| Tp0733 | outer membrane beta-barrel protein CDS | ggggacaagtttgtacaaaaaagcaggcttcATGAGCAGAACGTTC | ggggaccactttgtacaagaaagctgggtcCTAAAAGTGATAGCGC | WP_010882178.1 | 799993 | 800652 | 660 |
| Tp0750 | VWA domain-containing protein CDS | ggggacaagtttgtacaaaaaagcaggcttcATGCACCTGAAAAAAGCGC | ggggaccactttgtacaagaaagctgggtcCTAGTCAAGCGAAGGG | WP_010882195.1 | 816714 | 817385 | 672 |
| Tp0751 | vwb CDS | ggggacaagtttgtacaaaaaagcaggcttcGTGAATAGACCTTTACTGAGTGT | ggggaccactttgtacaagaaagctgggtcGGGCGAAGGAGCACTAGCCGGA | WP_010882196.1 | 817422 | 818135 | 714 |
| Tp0768 | membrane protein TmpA CDS | ggggacaagtttgtacaaaaaagcaggcttcATGAATGCTCATACGCTTGTG | ggggaccactttgtacaagaaagctgggtcTCGAGAGGCTCCTTCTTCG | WP_010882213.1 | 835524 | 836561 | 1038 |
| Tp0769 | LysM peptidoglycan-binding domain-containing protein | ggggacaagtttgtacaaaaaagcaggcttcATGAAGACACGTAATTTCTCGC | ggggaccactttgtacaagaaagctgggtcCGGCAGAGGACGGTATTTCACG | WP_010882214.1 | 36559 | 37537 | 978 |
| Tp0821 | MetQ/NlpA family ABC transporter substrate-binding protein CDS | ggggacaagtttgtacaaaaaagcaggcttcATGAAAGGAAAAACGGTGAGCGC | ggggaccactttgtacaagaaagctgggtcCAAAGCAGGCGCCACCTCC | WP_010882265.1 | 890493 | 891299 | 807 |
| Tp0859 | UPF0164 family protein CDS | ggggacaagtttgtacaaaaaagcaggcttcTTGGTGCGGCGGCCGTGTGTGTCG | ggggaccactttgtacaagaaagctgggtcCCCCCCCCCCCGGGTGTGGCGA | WP_014342835.1 | 937319 | 938803 | 1485 |
| Tp0865 | UPF0164 family protein CDS | ggggacaagtttgtacaaaaaagcaggcttcGTGGTACGGATGAGGCGGAG | ggggaccactttgtacaagaaagctgggtcGTGAAGAAAATGAAAATCG | WP_010882309.1 | 944967 | 946406 | 1440 |
| Tp0868 | flagellar filament 34.5 kDa core protein | ggggacaagtttgtacaaaaaagcaggcttcATGATTATCAATCACAACATG | ggggaccactttgtacaagaaagctgggtcCCGGAGAATTGAGAGAATCG | WP_010882311.1 | 948754 | 949614 | 861 |
| Tp0870 | FlaB3 | ggggacaagtttgtacaaaaaagcaggcttcATGATTATCAATCACAACATG | ggggaccactttgtacaagaaagctgggtcCTGCATCAAGCGGAGCACTGCC | WP_010882313.1 | 949893 | 950750 | 858 |
| Tp0965 | efflux RND transporter periplasmic adaptor subunit CDS | ggggacaagtttgtacaaaaaagcaggcttcATGACCACAGCACAGAAACTCC | ggggaccactttgtacaagaaagctgggtcTTTCCTCGCTGCACTTTGGTC | WP_010882409.1 | 1048161 | 1049123 | 963 |
| Tp0966 | hypothetical protein CDS | ggggacaagtttgtacaaaaaagcaggcttcGTGATCGCACGCAGGATGC | ggggaccactttgtacaagaaagctgggtcGAGTACTGGTCTGAACGAAGACGC | WP_010882410.1 | 1049148 | 1050782 | 1635 |
| Tp0967 | hypothetical protein CDS | ggggacaagtttgtacaaaaaagcaggcttcGTGAGCGGTCCGCGGGGTGG | ggggaccactttgtacaagaaagctgggtcTGTTTGGGTTCCAGAAG | WP_010882411.1 | 1050829 | 1052382 | 1554 |
| Tp0968 | hypothetical protein CDS | ggggacaagtttgtacaaaaaagcaggcttcGTGAGCGTAAGTTATCGTGGC | ggggaccactttgtacaagaaagctgggtcCGGCGCCACCTGTGGCACCAGG | WP_010882412.1 | 1052379 | 1054001 | 1623 |
| Tp0969 | hypothetical protein CDS | ggggacaagtttgtacaaaaaagcaggcttcTTGAAAGTTCTCCTGCGCG | ggggaccactttgtacaagaaagctgggtcCGAAGACCTCTCTTTCTTTTC | WP_014342850.1 | 1053998 | 1055572 | 1575 |
| Tp0971 | iron transporter CDS | ggggacaagtttgtacaaaaaagcaggcttcATGAAGAGGGTGAGTTTGCTCG | ggggaccactttgtacaagaaagctgggtcCCACTGAGGCCCCTTCCATTC | WP_010882415.1 | 1055752 | 1056366 | 615 |
| Tp1016 | basic membrane protein CDS | ggggacaagtttgtacaaaaaagcaggcttcATGGGCAGATACATAGTTCCCGC | ggggaccactttgtacaagaaagctgggtcCCAGTCGAGCACCTTGCCGAGC | WP_010882460.1 | 1108399 | 1109484 | 1086 |

^a^*attB* adapter sequence is in lowercase, *Tp*-locus specific sequence is in uppercase

Supplementary Table 2. Synthetic DNA cloned into expression plasmid pDEST203

| *Tp* locus | other name(s) | NCBI ID | Minimum | Maximum | Length (nt) |
| --- | --- | --- | --- | --- | --- |
| Tp0009 | Treponema pallidum repeat (Tpr) protein A (Msp) | H71379 | 8343 | 10165 | 1822 |
| Tp0061 | rpsR CDS | WP_010881510.1 | 69756 | 70055 | 300 |
| Tp0076 | carbohydrate ABC transporter permease CDS | WP_014342258.1 | 83400 | 84230 | 831 |
| Tp0119 | ABC transporter permease CDS | WP_010881568.1 | 138151 | 138810 | 660 |
| Tp0136 | hypothetical protein CDS | WP_010881584.1 | 157943 | 159430 | 1488 |
| Tp0163 | troA CDS, Tromp1 | WP_010881610.1 | 185763 | 186689 | 927 |
| Tp0177 | hypothetical protein (basic) | WP_252507609.1 | 194305 | 195612 | 1307 |
| Tp0214 | hypothetical protein CDS | WP_010881662.1 | 218717 | 218914 | 198 |
| Tp0216 | dnaK CDS | WP_010881664.1 | 219898 | 221805 | 1908 |
| Tp0248 | Uncharacterized lipoprotein | O83276.1 | 260962 | 261364 | 402 |
| Tp0292 | OmpA family protein CDS | WP_010881741.1 | 306712 | 307965 | 1254 |
| Tp0313 | hypothetical protein CDS, TprE | WP_010881762.1 | 329144 | 331432 | 2289 |
| Tp0324 | translocation/assembly module TamB domain-containing protein CDS | WP_014342370.1 | 341047 | 345453 | 4407 |
| Tp0403 | fliJ CDS | WP_010881851.1 | 429567 | 430019 | 453 |
| Tp0421 | tetratricopeptide repeat protein CDS | WP_010881869.1 | 448997 | 451048 | 2052 |
| Tp0433 | acidic repeat protein CDS | WP_014342796.1 | 461695 | 463509 | 1815 |
| Tp0479 | DUF2715 domain-containing protein CDS | WP_010881928.1 | 510487 | 511161 | 675 |
| Tp0515 | LPS-assembly protein LptD CDS | WP_010881964.1 | 554924 | 557899 | 2976 |
| Tp0558 | nickel/cobalt transporter CDS | WP_010882005.1 | 605758 | 606666 | 909 |
| Tp0625 | hypothetical protein CDS | WP_010882071.1 | 681849 | 682601 | 753 |
| Tp0654 | ABC transporter permease CDS | WP_010882099.1 | 719339 | 720157 | 819 |
| Tp0693 | hypothetical protein CDS | WP_010882138.1 | 761812 | 763134 | 1323 |
| Tp0718 | fliP CDS | WP_010882163.1 | 787485 | 788300 | 816 |
| Tp0728 | flgD CDS | WP_010882173.1 | 795008 | 795469 | 462 |
| Tp0772 | hypothetical protein CDS | WP_010882217.1 | 841335 | 842159 | 825 |
| Tp0792 | flagellin CDS | WP_010882237.1 | 859694 | 860554 | 861 |
| Tp0855 | hypothetical protein CDS | WP_010882299.1 | 931186 | 934569 | 3384 |
| Tp0856 | UPF0164 family protein CDS | WP_014342833.1 | 934725 | 935909 | 1185 |
| Tp0858 | UPF0164 family protein CDS | WP_010882302.1 | 936011 | 937237 | 1227 |
| Tp0897 | TprK | AF194369.1 | 135 | 1647 | 1512 |
| Tp0923 | PEGA domain-containing protein CDS | WP_014342645.1 | 1003265 | 1004287 | 1023 |
| Tp0951 | rpmH CDS | WP_010882395.1 | 1034084 | 1034239 | 156 |
| Tp0954 | tetratricopeptide repeat protein CDS | WP_010882398.1 | 1036755 | 1038191 | 1437 |
| Tp0993 | septal ring lytic transglycosylase RlpA family protein CDS | WP_014342852.1 | 1078854 | 1079879 | 1026 |
| Tp1031 | hypothetical protein CDS, TprL | WP_010882475.1 | 1125976 | 1127520 | 1545 |
| Tp1038 | DNA starvation/stationary phase protection protein CDS, TpF1, 4D | WP_010882482.1 | 1136431 | 1136964 | 534 |

Supplementary Table 3. Other expression plasmids

| Tp locus | Protein Name | NCBI ID | Vector | Insert (AA) | Reference (PMID) |
| --- | --- | --- | --- | --- | --- |
| Tp0326 | bamA CDS, Tp92 | WP_014342788.1 | pET26b | 22-837 | 25825429 |
| Tp0327 | OmpH family outer membrane protein CDS | WP_010881775.1 | pET28a | 23-172 | 32238570 |
| Tp0369 | bamD CDS | WP_010881817.1 | pDEST17 | 16-516 |  |
| Tp0486 | hypothetical protein CDS | WP_012460559.1 | pDEST17 | 19-531 |  |
| Tp0557 | DUF1007 family protein CDS | WP_010882004.1 | pRSETc | 26-237 | 16239550 |
| Tp0620 | Tpr protein I CDS | WP_014342810.1 | pET23b | 18-609 | 25805501 |
| Tp0621 | Tpr protein J CDS | WP_010882067.1 | pRSET/C | 269-494 | 17030565 |
| Tp0622 | tetratricopeptide repeat protein CDS | WP_010882068.1 | pETHisTEVN-term | 30-589 |  |
| Tp0624 | OmpA family protein CDS | WP_010882070.1 | pETHisTEVN-term | 61-476 | 27832149 |
| Tp0648 | tetratricopeptide repeat protein CDS | WP_010882093.1 | pETHisTEVN-term | 1-682 |  |
| Tp0742 | Obg family GTPase CgtA CDS | WP_010882187.1 | pDEST17 | 1-376 |  |
| Tp0773 | trypsin-like peptidase domain-containing protein CDS | WP_014342572.1 | pET28a | 20-398 |  |
| Tp0784 | hypothetical protein CDS | WP_010882229.1 | pET28a | 18-199 |  |
| Tp0785 | hypothetical protein CDS | WP_010882230.1 | pET28a | 1-163 |  |

Supplementary Table 4. Labeled antibodies for flow cytometry

| Assay | Antibody | Clone | Vendor |
| --- | --- | --- | --- |
| AIM/sort | anti-CD3-PE | UCHT1 | Life Technologies |
|  | anti-CD8α-FITC | 3B5 | Life Technologies |
|  | anti-CD4-allophycocyanin-H7 | RPA-T4 | BD |
|  | anti-CD69-Brilliant Violet-421 | FN50 | Biolegend |
|  | anti-CD137-APC | 4B4-1 | BD |
| ICS | anti-CD49d | 9F10 | Biolegend |
|  | anti-CD28 | CD28.2 | Biolegend |
|  | anti-CD3-ECD | UCHT1 | Beckton Coulter |
|  | anti-CD4-FITC | RPA-T4 | Biolegend |
|  | anti-CD8-PerCP-Cy5.5 | SK1 | BD |
|  | anti-IFN-γ-PE | 45.B3 | BD |
|  | anti-IL-2-APC | MQ1-17H12 | BD |

Supplementary Table 5. Please see file “Reid_Tp-specific_Tcell_antigens_Sup_Materials_Table_5”

**A**

**B**

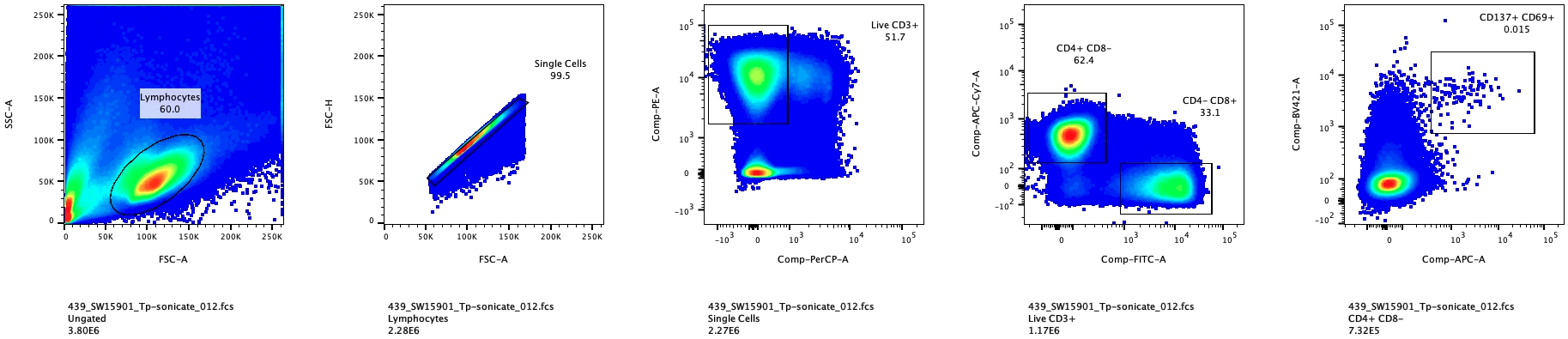

**Dead-PerCP**

**CD3-PE**

**CD8-FITC**

**CD4-APC-Cy7**

**FSC-A**

**FSC-H**

**FSC-A**

**SSC-A**

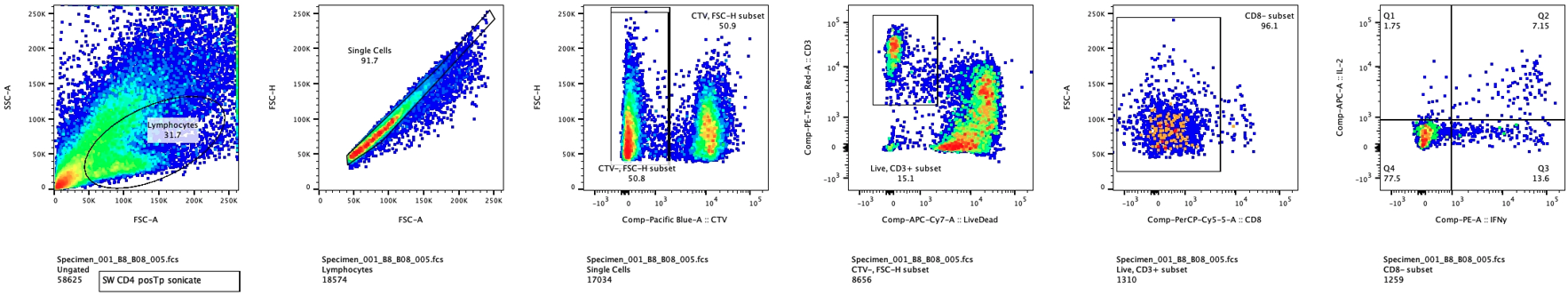

**FSC-A**

**SSC-A**

**FSC-A**

**FSC-H**

**FSC-H**

**PBMC-CTV**

**CD3-PE**

**Dead-APC-CY7**

**FSC-A**

**CD8-PerCP-Cy5.5**

Supplementary Figure 1. Representative gating scheme prior to Activation Induced Marker (AIM) sorting or prior to final analysis of intracellular cytokine staining (ICS) assays. *A*, Gating used for PBMC AIM, showing lymphocytes, single cells, CD3+ live cells, then CD4+ and CD8- cells. *B,* Gating used for ICS, isolating from left to right lymphocytes, single cells, the expanded responder T cell line, CD3+ live cells, then CD8- cells.

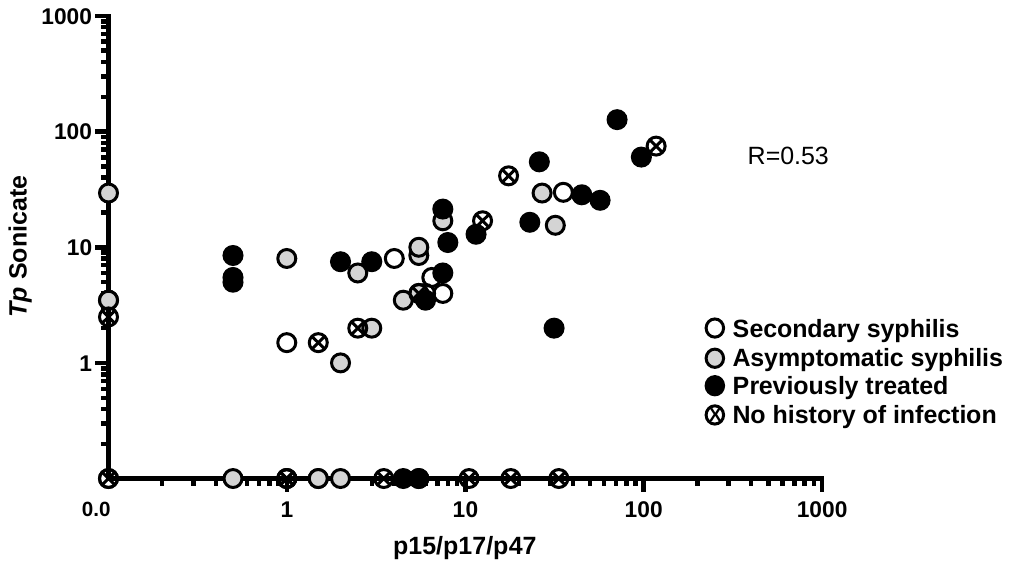

Supplementary Figure 2. Correlation between PBMC IFNγ responses to *Tp* sonicate and recombinant p15/p17/p47 antigen. Values are means of background-subtracted duplicate ELISPOT wells. Specimens from participants are indicated per clinical status.

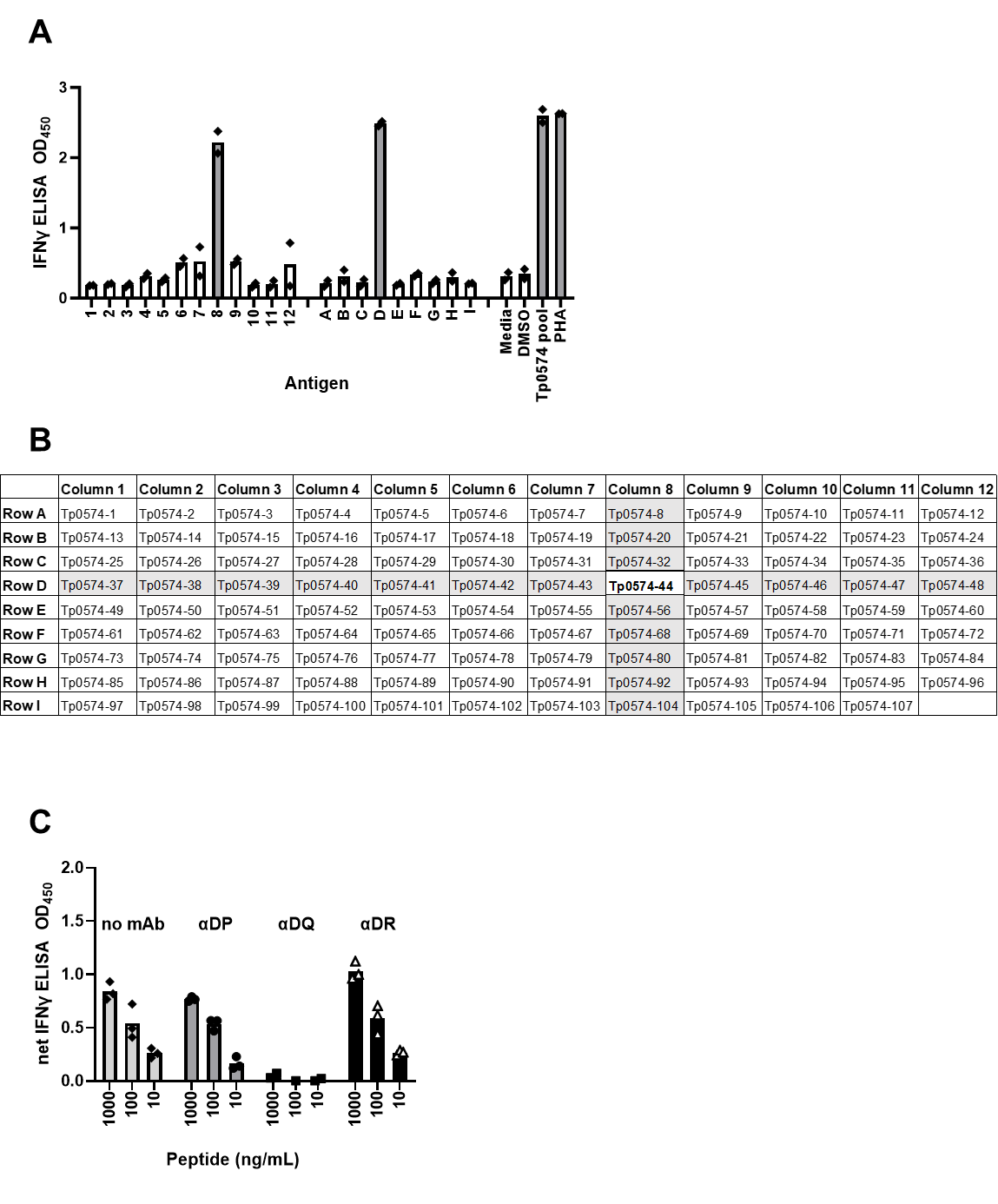

Supplementary Figure 3. CD4 T cell epitope mapping and determination of HLA restriction to the locus level.

Representative data from participant 05, treated for syphilis of unknown duration 120 months before enrollment. *A,* Tp0574 epitope mapping using pooled peptides. Individual peptide final concentrations are 1 μg/mL.  Reactivity of polyclonal *Tp* -reactive CD4 T cells detected by IFNγ secretion using autologous EBV-LCL as APC.  Graph shows means (bars) of technical duplicates (dots). Controls include a pool of all 107 peptides in Tp0574. *B,* Peptide Tp0574-44 (AA 173-185) is at the overlap between active column pool 8 and row pool D. *C,* Graph shows means (bars) of technical triplicates (shapes). Reactivity to peptide Tp0574 (AA 173-185) is confirmed at left in the absence of anti-HLA class II blocking antibody.  Anti-HLA-DQ completely inhibited IFNγ response to Tp0574 AA 173-185 while antibodies to HLA-DR or-DP had no effect.

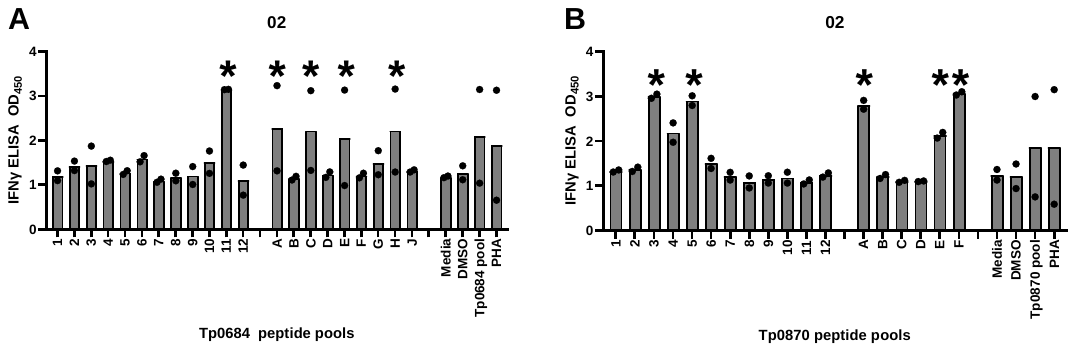

Supplemental Figure 4. TCL IFNγ secretion in response to peptides and epitope mapping.

Epitope mapping using matrix pools of 6-12 peptides where individual peptide final concentrations are 1 μg/mL. Reactivity of polyclonal *Tp*-reactive CD4 T cells detected by IFNγ secretion using autologous EBV-LCL as APC. Graph shows means (bars) of technical duplicates (dots). *A,* 4 peptides within reactive pool 11 were further assessed for individual peptide reactivity. *B*, Peptides common to pools 3, 5, A, E and F were further assessed individually. Controls at right include media, DMSO, *Tp* sonicate (*Tp*), pool of all peptides covering a given open reading frame, and PHA. *Pools were selected (both replicate responses ≥ 2x DMSO response) to deduce reactive peptides for individual evaluation.

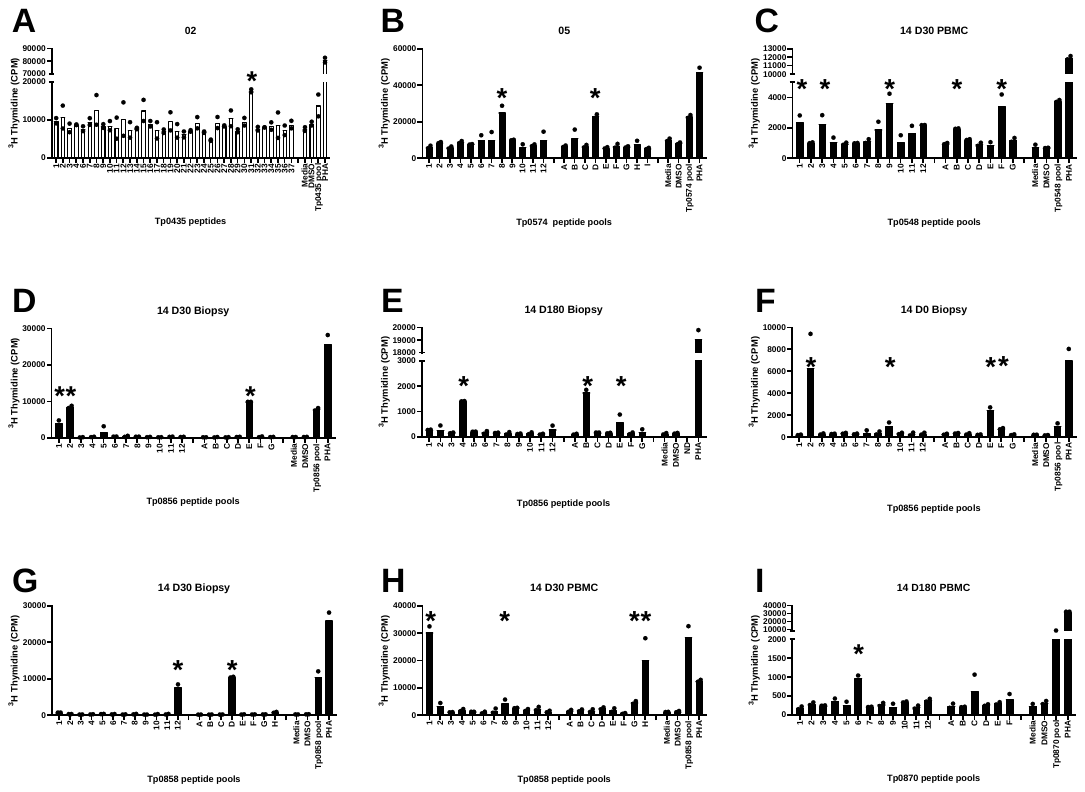

Supplemental Figure 5. TCL proliferation in response to peptides and epitope mapping.

Epitope mapping using matrix pools of 6-12 peptides except for Tp0435 (*A*) where individual peptides were used. Individual peptide final concentrations are 1 μg/mL. Reactivity of polyclonal *Tp* -reactive CD4 T cells detected proliferation expressed as counts per minute (CPM) using autologous EBV-LCL as APC. Graph shows means (bars) of technical duplicates (dots). Controls at right include media, DMSO, *Tp* sonicate (*Tp*), pool of all peptides covering a given open reading frame, and PHA. ND = not done. *pools and peptides selected (both replicate responses ≥ 2xDMSO response) to deduce reactive peptides for individual evaluation.
