## Supplementary Methods for "*Treponema pallidum* periplasmic and membrane proteins are recognized by circulating and skin CD4+ T cells"

**Antigens**

*Proteins*

Recombinant proteins were expressed as follows: DNA was extracted from *Tp* Nichols strain lysate prepared from infected rabbit testicles using the QIAmp DNA Mini Kit (QIAGEN). PCR primers with recombinase-mediated integration sites were designed to amplify full-length or truncated open reading frames (ORFs) **(Supplementary Table 1)** using the Seattle Nichols genome (Genbank NC_CP010422). PCR reactions (50 µL) contained 300 µM dNTPs, 0.4 μM primers and 5 U LA Hotstart Taq DNA Polymerase (Takara Bio). Amplicons were cloned into pDONR221 using Gateway BP Clonase (ThermoFisher). Alternatively, *Tp* genes with *att*B flanking recombination sites were synthesized and inserted into pUC-GW-AMP (Azenta) **(Supplementary Table 2)**. The *Tp* amplicon sequences were confirmed by Sanger sequencing (Azenta) and subcloned into pDEST203 using Gateway LR Clonase II (ThermoFisher) [1]. Proteins were expressed using the Expressway™ in vitro transcription/translation (IVTT) system (ThermoFisher) with an amino terminal histidine tag, except for Tp0326 which was expressed with a carboxy terminal histidine tag. *P. falciparum* proteins, expressed similarly, were used as controls. Expression of molecular clones was confirmed by ani-6x-histidine dot blot (not shown).

**Enrichment and expansion of *Tp*-specific T cells**

Activation-induced marker (AIM) based sorting allows for recovery of antigen reactive cells for downstream studies. PBMC (4x10^6^) were incubated with *Tp* or mock sonicate (1/10 dilution) for 18h at 37°C, 5% CO_2_ in T cell medium (TCM; RPMI-1640, 4% Human Serum (Valley Biomedical), 4% FBS and 2 mM L-glutamine, 50 U/mL penicillin/streptomycin). Before (AIM)-based cell sorting, cells were labeled with antibodies listed in **Supplementary Table 4** for 30 minutes at room temperature in 50 μL TCM, washed and resuspended (1x10^7^ cells/mL) in TCM. Viability was assessed by 7-AAD addition. CD4+ T cells that expressed CD69 and CD137 after 18 hour exposure of PBMC to *Tp* antigen were isolated by sorting **(Figure 1B and Supplementary Figure 1A)**.

For skin biopsies, tissue was minced in a small volume of TCM and fragments (5-10 per well) used for T cell isolation as described [2].T cells from skin biopsy fragments, and candidate *Tp*-specific PBMC sorted as D3+CD4+CD69+CD137+ cells, were non-specifically expanded in culture with 10^6^/mL γ-irradiated (3,300 rads) allogeneic PBMC and PHA-P (1.6 μg/mL) in flat bottom 48 well plates and 98 well round bottom plates, respectively. Natural human IL-2 (32 U/mL, Hemagen) was added at 48 hours and maintained for 14–16 days. A portion of the cells from the initial expansion from blood or skin were further polyclonally expanded with anti-CD3 (clone OKT3, Biolegend) with addition of recombinant IL-2 starting at day four, for 14-16 days [2].

The reactivity of expanded *Tp*-specific T cell lines (TCL) was assessed by intracellular cytokine staining (ICS) as described [1]. Briefly, TCL added to an equal number of labeled autologous PBMC [CellTrace Violet (CTV, ThermoFisher)] were stimulated with *Tp* or mock sonicates (1/10 dilution) for 16 h in TCM containing anti-CD49d and anti-CD28 (Biolegend); brefeldin A was added at two hours. Negative and positive controls included medium, and PHA-P. Cells were fixed (BD FACS Lysing Solution), permeabilized (BD FACS Permeabilizing Solution 2), and stained with the labeled antibodies listed in **Supplementary Table 4**. Data collected on an Aurora cytometer (Cytek) was analyzed with FlowJo 10 (BD). Responder TLC cells were gated away from CTV-labeled PBMC for analysis as described [1] **(Supplementary Figure 1B)**. We measured enrichment in expanded TCL by the ratio of net ICS positive cells (expressing IFNγ, IL-2 or both cytokines in response to *Tp*) to the net AIM response to *Tp* sonicate.
